# Auxin-degron system identifies immediate mechanisms of Oct4

**DOI:** 10.1101/2020.09.21.306241

**Authors:** Lawrence E Bates, Mariana R P Alves, José C R Silva

## Abstract

The pluripotency factor Oct4 is essential for the maintenance of naïve pluripotent stem cells in vitro and in vivo. However, the specific role of Oct4 in this process remains unknown. Here, we developed a rapid protein-level Oct4 depletion system that demonstrates that the immediate downstream response to loss of Oct4 is reduced expression of key pluripotency factors. Our data show a requirement for Oct4 for the efficient transcription of several key pluripotency factors, and suggest that expression of trophectoderm markers is a subsequent event. Additionally, we find that Nanog is competent to bind to the genome in the absence of Oct4, and this binding is in fact enhanced. Globally, however, active enhancer associated histone mark H3K27ac is depleted. Our work establishes that while Oct4 is required for the maintenance of the naïve transcription factor network, at a normal ESC level it antagonises this network through inhibition of Nanog binding.

## Introduction

Naïve pluripotent stem cells (nPSCs) are the embryonic founders of the cells present in the adult animal. The transcription factor Oct4, expressed from the gene Pou5f1, is necessary for the maintenance of naïve pluripotency in vivo and in vitro. In both cases, loss of Oct4 ultimately leads to exit from naïve pluripotency and with cells taking on characteristics of the trophoblast lineage, including expression of marker genes Cdx2, Pl-1 and Eomes, and adoption of trophectoderm-like morphology (Nichols et al., 1998; Niwa et al., 2000).

Despite this, it is not understood why Oct4 is essential to the naïve state; Oct4 is a highly promiscuous transcription factor, binding to thousands of sites in the genome (King and Klose, 2017; Simandi et al., 2016), and many of its target genes are known not to be essential for the maintenance of nPSCs (Hall et al., 2009; Matoba et al., 2006; Zwaka, 2012). At the same time, no target has been identified that can rescue Oct4-loss driven differentiation either through overexpression or knockout (Hall et al., 2009; Matoba et al., 2006). It therefore seems likely that differentiation upon loss of Oct4 is the result of misregulation of a number of genes rather than of any single critical target.

Studying the effects of loss of Oct4 are further complicated by the fact that Oct4 appears to function in a level-dependent manner (Niwa et al., 2000). Mouse nPSCs engineered to express reduced levels of Oct4 are recalcitrant to differentiation and in some cases can be maintained in minimal culture conditions (Karwacki-Neisius et al., 2013; Radzisheuskaya et al., 2013), including basal serum-free media devoid of otherwise essential growth factors and inhibitors of differentiation pathways. Expression of pluripotency associated genes may be enhanced in this state, and responses to signalling cues may be altered. In particular, it appears that transcript and protein levels of the core pluripotency factor Nanog are negatively correlated with Oct4 expression (Karwacki-Neisius et al., 2013).

Nanog is a homeodomain transcription factor that co-binds with Oct4 at many enhancers to promote expression of other pluripotency genes (Chen et al., 2008; Loh et al., 2006). While Nanog is not absolutely required for maintenance of the naïve identity (Chambers et al., 2007), loss of Nanog results in widespread differentiation and reduced self-renewal capacity (Chambers et al., 2007; Ivanova et al., 2006; Mitsui et al., 2003). Interestingly, binding of Nanog to the genome appears to be enhanced in self-renewing nPSCs expressing low levels of Oct4, in line with the increased overall level of Nanog protein (Karwacki-Neisius et al., 2013; Radzisheuskaya et al., 2013), while during differentiation induced by loss of Oct4 Nanog binding is reduced (King and Klose, 2017). However, the immediate effect of total loss of Oct4 protein on Nanog binding to the genome remains unknown.

Conventional methods for depleting Oct4, such as genetic ablation via Lox-Cre systems or transcriptional repression using Tet-OFF regulated transgenes, rely on natural degradation to remove Oct4 mRNA and protein. A result of this is that Oct4 persists for a long time. Consequently, responses to depletion of Oct4 are typically examined on the order of hours to days following manipulation (Hall et al., 2009; Niwa et al., 2000), making it impossible to discern whether results are related to Oct4 or simply to cells undergoing differentiation. Further complicating such analysis, microarray data have revealed changes in the expression of thousands of genes at early timepoints after Oct4 suppression, even as the majority of Oct4 protein remains (Hall et al., 2009). This means that such variation may represent a response to a change in Oct4 expression to lower than WT levels, previously described to enhance self-renewal capacity in nPSCs (Karwacki-Neisius et al., 2013; Radzisheuskaya et al., 2013). Thus, the nature of the essential role of Oct4 for the self-renewal of nPSCs remains elusive.

In order to overcome these confounding factors, we generated an Oct4 fusion protein to an auxin-inducible degron. This allows us to induce rapid protein-level depletion of Oct4 making it possible for the first time to study the immediate molecular responses to loss of Oct4. These revealed an unprecedented impact on the transcriptional activity of pluripotency-associated transcription factor genes and addressed a long-standing question regarding the requirement of Oct4 for the binding of a key pluripotency factor, Nanog, to regulatory sequences.

## Results

### Auxin-degron tagged Oct4 sustains naïve PSC self-renewal and permits rapid loss of Oct4

Oct4 protein has a relatively long half-life; unlike pluripotency factor Nanog with a reported half-live of around 3 hours, the half-life of Oct4 protein is typically found to be >6 hours to as much as 24 hours (Abranches et al., 2013; Muñoz Descalzo et al., 2013). Additionally, the endogenous Oct4 mRNA also appears to be unusually stable in mouse ESCs (Abranches et al., 2013). Indeed, we found that conventional tamoxifen-induced CreER driven genetic ablation of Oct4 resulted in a gradual reduction, and required over a day to fully deplete Oct4 RNA and protein (Figure 1A-B). Four days after induction, cells showed distinct morphological changes, resembling trophoblasts and more differentiated trophectodermal cells (Figure S1A). This shows that genetic and transcriptional methods of silencing Oct4 take a long time to fully deplete Oct4 protein, and cells are likely to experience a protracted period of low Oct4 content. Given the transcriptional and phenotypic effects previously observed in cells stably expressing low levels of Oct4 (Karwacki-Neisius et al., 2013; Radzisheuskaya et al., 2013), this is likely to have confounding effects on studies of immediate responses to complete removal of Oct4. We therefore decided to utilize an auxin-inducible degron (AID) tag (Baker et al., 2016; Nishimura et al., 2009) to achieve rapid, inducible protein-level degradation of Oct4 on addition of the small molecule IAA.

**Figure 1.**
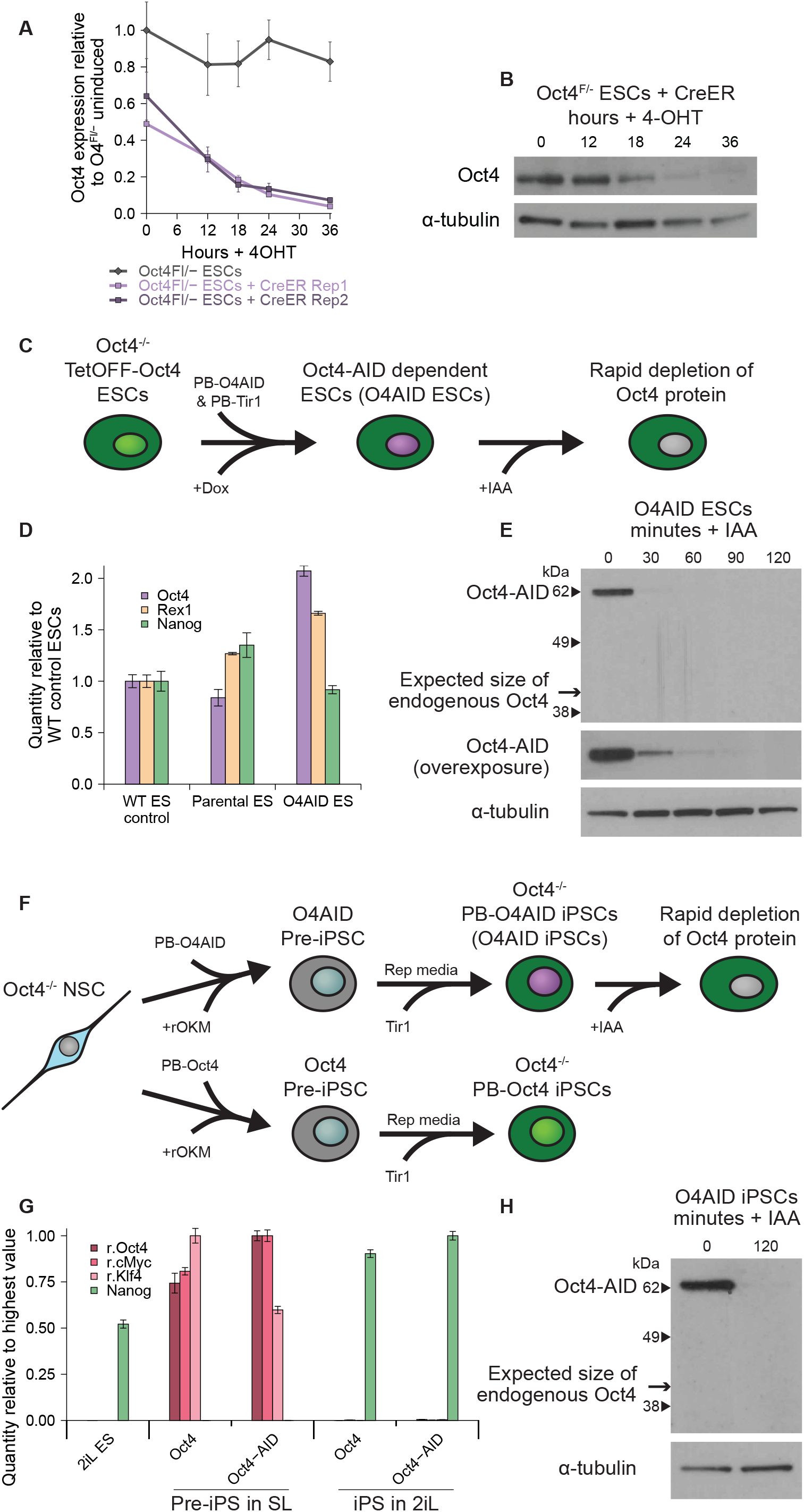
Auxin-degron tagged Oct4 sustains naïve PSC self-renewal and permits rapid loss of Oct4. A&B) Kinetics of Oct4 depletion in conventional Oct4^Fl/-^ ESCs were examined. A) Oct4 expression level (RT-qPCR) following addition of 4-OHT and media change to SL. B) Oct4 protein level (Western blot) following addition of 4-OHT and media change to SL. α-tubulin shown as a loading control. C) Schematic showing the generation and use of O4AID ESCs. D) Expression profiling (RT-qPCR) of pluripotency markers. E) Oct4-AID fusion and wild-type Oct4 protein level (Western blot) following addition of IAA. α-tubulin loading control. F) Schematic showing the generation and use of O4AID iPSCs. rOKM, retroviral Oct4, Klf4 and cMyc. G) Expression profiling (RT-qPCR) of retroviruses and pluripotency factor Nanog in partially and fully reprogrammed cells. r.Oct4 – retroviral Oct4; r.cMyc – retroviral cMyc; r.Klf4 – retroviral Klf4. H) Oct4-AID fusion and wild-type protein level (Western blot) following addition of IAA. RT-qPCR data represent mean ± SD of three technical replicates.

Since Oct4 is essential for the maintenance of pluripotency we sought to confirm that AID-tagged Oct4 retained its biological function by testing its capacity to support ESCs lacking wild-type Oct4. We utilized an existing TetOFF-Oct4 cell line; non-functional transgenes cannot rescue self-renewal on dox-induced inhibition of wild-type Oct4 in these cells (Niwa et al., 2002), allowing them to be used in a complementation assay. We generated Oct4-/- TetOFF-Oct4 Oct4-AID ESCs (O4AID ESCs hereafter) by transfecting these cells with constitutively expressed Piggybac Oct4-AID and Tir1 constructs and maintained cells in the presence of dox to inhibit expression of wild-type Oct4 (Figure 1C). Cells continued to proliferate and showed normal morphology (Figure S1B). Cells expressed the fusion protein with no detectable wild-type Oct4 present, and did not show substantially altered expression of key pluripotency genes (Figure 1D-E). This demonstrates that the Oct4 fusion protein retains its original capacity to maintain naïve pluripotency and is not substantially altered in its function by the addition of the AID domain.

To further validate the function of Oct4 within the fusion protein and to establish a second independent cell system, we generated iPSCs null for endogenous Oct4 and therefore wholly reliant on either transgenic wild-type Oct4 or Oct4-AID for their maintenance (Figure 1F). After a period of outgrowth, cells initiated reprogramming and generated intermediate pre-iPSCs (Silva et al., 2008; Theunissen et al., 2011) with high expression of retroviral reprogramming factors but very low expression of the pluripotency factor Nanog (Figure 1G). Under naïve-specific conditions, colonies with a domed, iPS-like morphology were readily obtained using either wild-type or Oct4-AID constructs (Figure S1C). Nanog was robustly expressed while retroviral transgenes were efficiently silenced in the fully reprogrammed iPSCs (Figure 1G). Again, as expected wild-type Oct4 was not detectable while the Oct4-AID fusion protein was robustly expressed (Figure 1H). The ability of the Oct4-AID transgene to maintain pluripotency in this Oct4 knockout background further demonstrates that the essential function of Oct4 is maintained.

Utilizing the new protein degradation system, tagged Oct4 protein levels could be greatly reduced within half an hour and undetectable within two hours of addition of IAA (Figure 1E&H). This quick turnover means that there is no appreciable Oct4-low state, and we therefore sought to use this system to study immediate responses of cells to total loss of Oct4.

### Oct4 is required for the expression of key pluripotency factors

We analysed gene expression changes following conventional tamoxifen-induced CreER-driven genetic ablation of Oct4 (Figure 2A and Figure S2A). The results indicate that upregulation of differentiation markers, particularly trophoblast associated genes, occurs concurrently with the downregulation of some naïve pluripotency factors. In keeping with previous observations, however, many of these gene expression changes are detected while significant amounts of Oct4 protein are still present (Figure 1B). However, other pluripotency-associated genes remain expressed at high levels for an extended period of time; indeed, expression of Nanog and Tfcp2l1 appear to increase following removal of Oct4 (Figure 2A and Figure S2A). This is consistent with reports that expression of Nanog may be enhanced in cells with reduced levels of Oct4 (Karwacki-Neisius et al., 2013; Radzisheuskaya et al., 2013), supporting the notion that conventional methods of Oct4 depletion pass through a protracted Oct4-low state, complicating interpretation of the effects of removal of Oct4.

**Figure 2.**
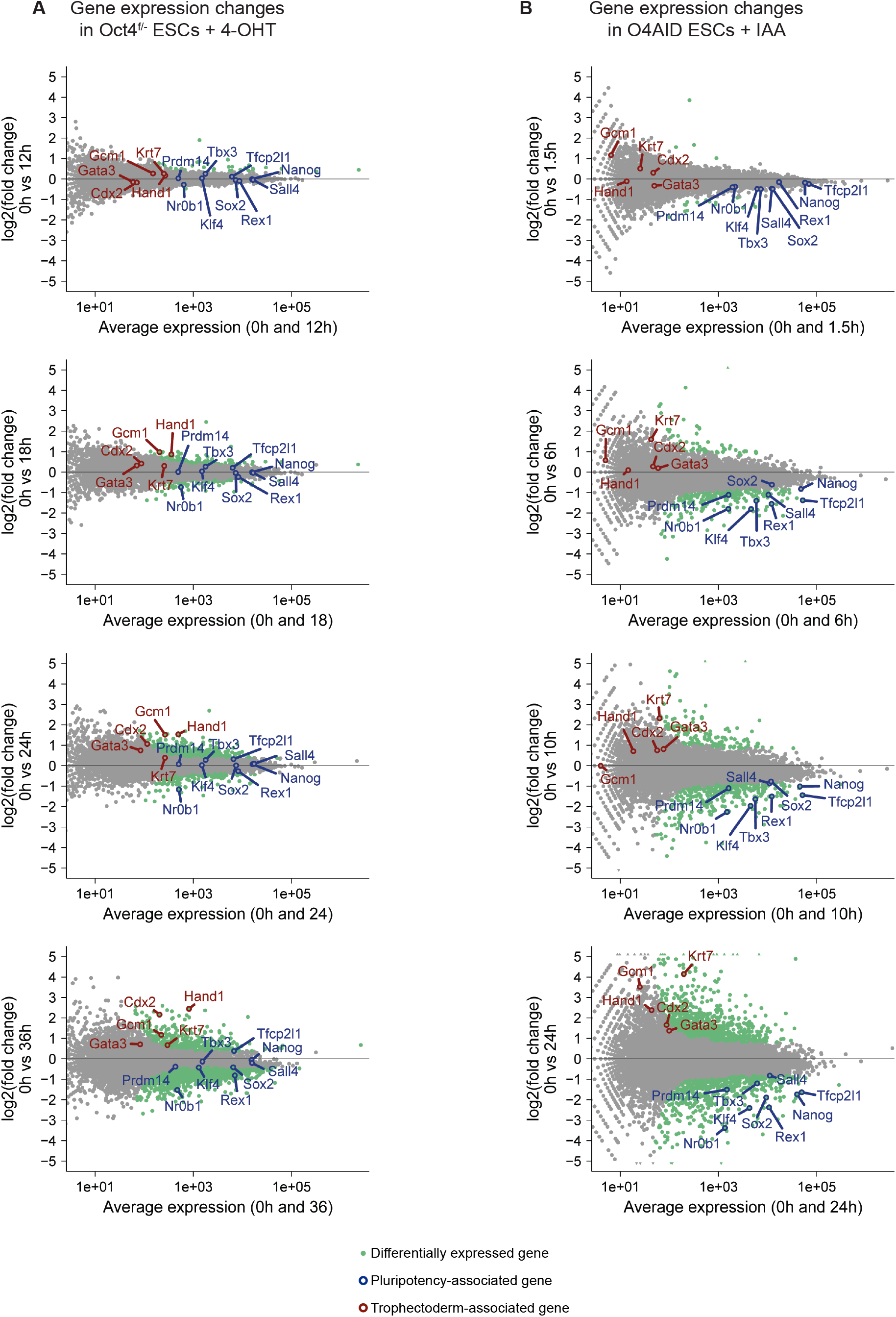
Oct4 is required for the expression of key pluripotency factors. MA plots showing gene expression (RNA-seq) changes following A) deletion of Oct4 in conventional Oct4^F/-^ CreER ESCs by addition of 4-OHT or B) degradation of Oct4 protein in O4AID ESCs by addition of IAA. Differentially expressed genes highlighted in green (q > 0.9, NOISeq-sim). Selected pluripotency (blue) and differentiation (red) associated genes indicated. A) top to bottom: 0 vs 12, 0 vs 18, 0 vs 24, and 0 vs 36 hours respectively. B) top to bottom: 0 vs 1.5, 0 vs 6, 0 vs 10, and 0 vs 24 hours respectively.

In contrast to the above, analysis of O4AID cells indicated that expression of naïve-associated factors is quickly extinguished on loss of Oct4 (Figure 2B and Figure S2B-C). Factors such as Tfcp2l1 and Nanog were rapidly downregulated following targeted Oct4 protein depletion whereas they were actively maintained in the slower conventional system. This highlights that there are important differences in transcriptional responses between gradually reducing Oct4 protein levels and removing Oct4 protein entirely.

Examining several trophoblast-associated genes confirmed that these cells exit pluripotency following loss of Oct4 and differentiate towards trophectodermal lineages (Figure 2B and Figure S2B-C), in keeping with conventional Oct4 depletion systems. However, strong upregulation of trophectoderm associated genes is only seen after a decrease in pluripotency marker expression using this system, indicating that the decision to enter this extraembryonic identity is made only after cells have begun to exit the naïve state.

### Enhancers show rapid epigenetic and functional inactivation following loss of Oct4

Given the rapid downregulation of pluripotency-associated genes, we examined how enhancer elements were affected by loss of Oct4 at a number of these loci. Enhancer-associated transcription can be observed at many active enhancers (Kim et al., 2010; De Santa et al., 2010), and can play a functional role in promoting transcription of target genes (Alvarez-Dominguez et al., 2017; Shii et al., 2017). We observed loss of transcription from several pluripotency enhancer elements shortly after ablation of Oct4 (Figure 3A), implying loss of enhancer activity.

**Figure 3.**
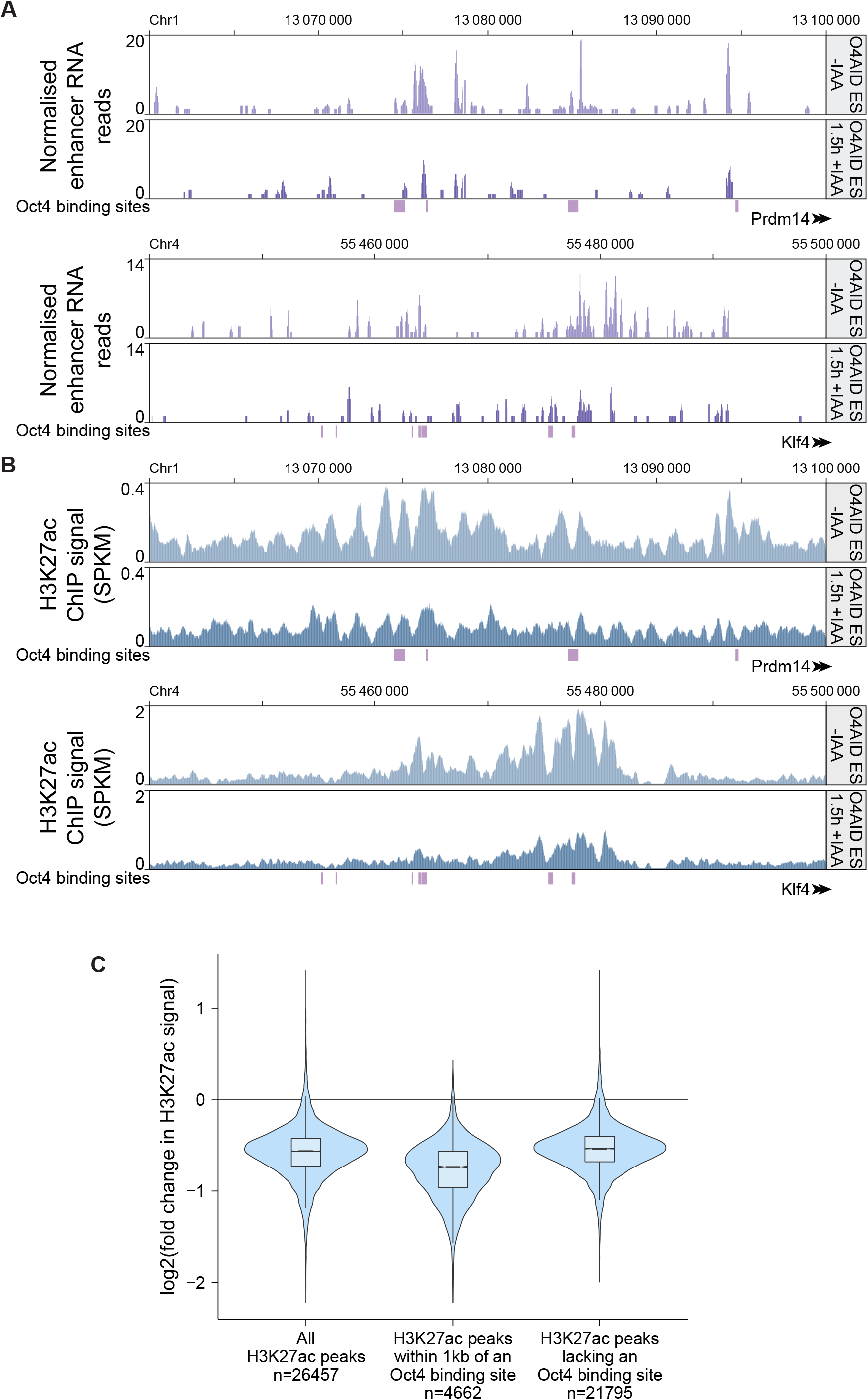
Oct4 is required for enhancer activity at key pluripotency loci, and for maintaining global H3K27ac. A) Mapped RNA-seq reads of enhancer RNAs at the Prdm14 distal enhancer (top) and the Klf4 distal enhancer (bottom) in O4AID ESCs before and 1.5 hours after addition of IAA. B) ChIP-seq analysis of H3K27ac at the Prdm14 distal enhancer (top) and the Klf4 distal enhancer (bottom) in O4AID ESCs before and 1.5 hours after addition of IAA. Genomic coordinates refer to the GRCm38/mm10 genome assembly, and gene intron/exon annotations are taken from Ensembl. Oct4 binding sites generated from ChIP-seq data from Marson et al. 2008 indicated in purple. C) Violin and box plot showing log2-fold change in H3K27ac signal between uninduced and 1.5 hour IAA treated O4AID ESCs. Boxes show the median value and extend to the 25^th^ and 75^th^ quartiles, and whiskers extend to 1.5 times the interquartile range. Plotted are all H3K27ac peaks (n=26457) above a (background) threshold, and the complementary subsets of peaks within 1kb of an Oct4 binding site (n=4662) and not within 1kb of an Oct4 binding site (n=21795). For each set a paired t-test of H3K27ac signal before and after treatment showed highly significant change, p < 10^−10^.

To validate this, we looked at the level of acetylation of histone H3 at lysine residue 27 (H3K27ac), a mark closely associated with active enhancers. Using ChIP-seq, we observed a dramatic decrease in H3K27ac at pluripotency-associated enhancers (Figure 3B and Figure S3A), and we confirmed this using ChIP-qPCR in both O4AID ES and O4AID iPS systems (Figure S3B-C). Loss of H3K27ac was not restricted to pluripotency-associated loci, however; global analysis of all H3K27ac peaks showed a significant decrease (Figure 3C). Using peaks called from Oct4 ChIP-seq data generated by (Marson et al., 2008), we examined if this was the result of decommissioning of Oct4-cobound enhancers; while the reduction in H3K27ac signal was stronger at sites close to Oct4 binding sites, H3K27ac peaks that were not directly bound by Oct4 still showed a marked reduction, indicating a remarkable shift in the global chromatin landscape in the absence of Oct4.

### Oct4 is dispensable for Nanog binding to pluripotent regulatory sequences

Depletion of Oct4 was so rapid that protein levels of other pluripotency factors were essentially unchanged by the time Oct4 protein was fully removed (Figure 4A and Figure S4A). As a result, we decided to use this opportunity to examine changes in the chromatin binding profile of Nanog, a transcription factor that binds to many of the same enhancer elements as Oct4.

**Figure 4.**
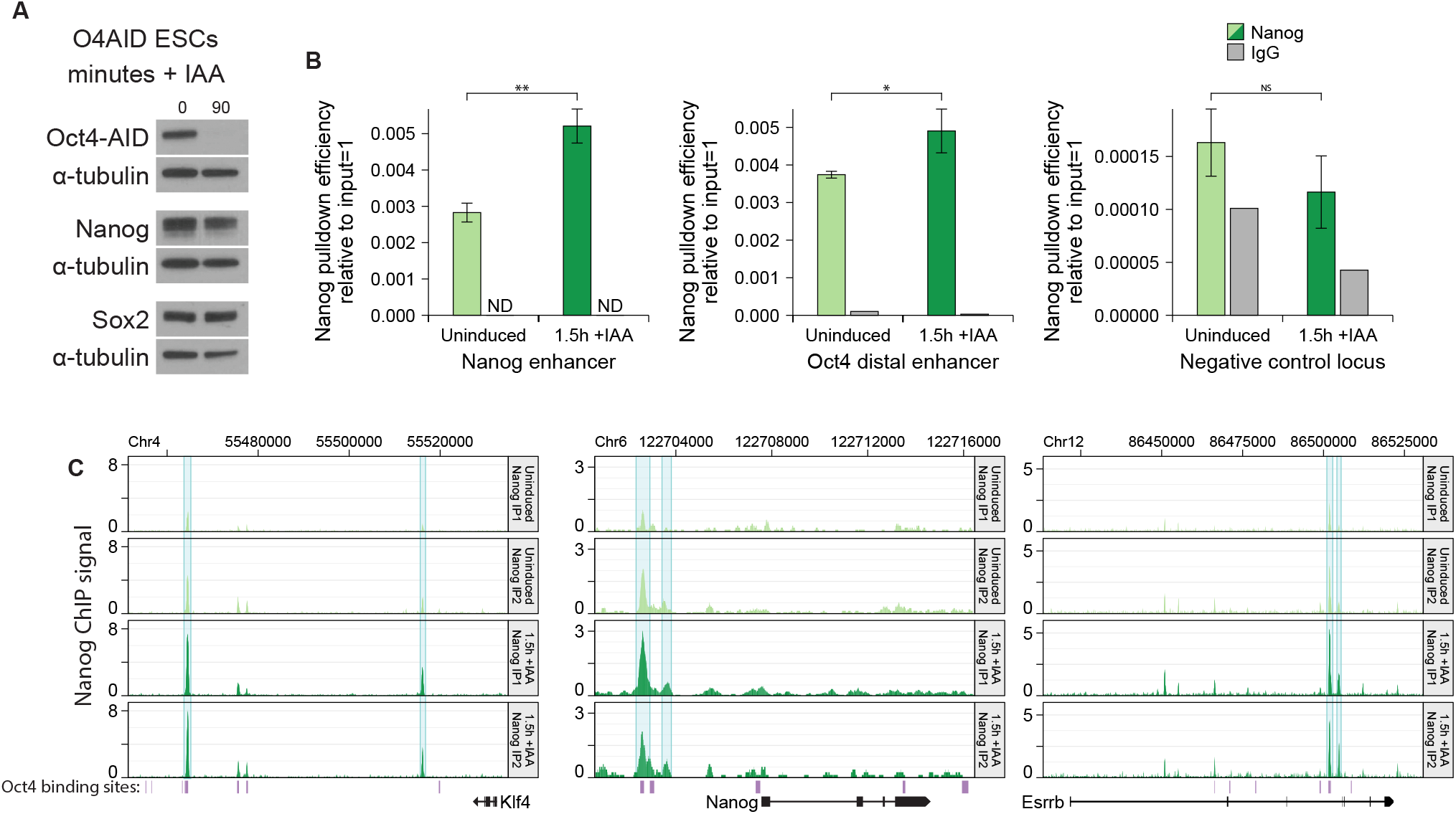
Oct4 is dispensable for Nanog binding to pluripotent regulatory sequences. A) Protein level of Oct4 and Nanog (Western blot) in O4AID ESCs before and 1.5 hours after addition of IAA with α-tubulin as a loading control. B) ChIP qPCR following pulldown of Nanog or using normal IgG negative control at Nanog binding sites or a negative control locus in O4AID ESCs. ChIP qPCR data for Nanog pulldown represent the mean ± SD of three IPs; IgG pulldown represents the mean of three technical replicates of single IP. ND = not detected. * P<0.05, ** P<0.005 (Student’s t test, two-tailed, unpaired, assuming equal variance). C) Visualization of Nanog ChIP-seq signal (SPKM) across indicated loci before and 1.5 hours after addition of IAA, shown for two IPs. Genomic coordinates refer to the GRCm38/mm10 genome assembly, and gene intron/exon annotations are taken from Ensembl. Oct4 binding sites generated from ChIP-seq data from Marson et al. 2008 indicated in purple.

Nanog binding efficiency at the enhancer of the Nanog and Oct4 genes was analysed by ChIP-qPCR. Surprisingly in both O4AID ES and iPS systems, Nanog not only remained bound but there was greater Nanog signal following depletion of Oct4 (Figure 4B and Figure S4B). Interestingly, other work has suggested that in the Oct4-low state Nanog has increased genomic occupancy, though Oct4 was still present (Karwacki-Neisius et al., 2013; Radzisheuskaya et al., 2013). In the data presented here we actively avoid the presence of Oct4, suggesting that either directly or indirectly Oct4 actively reduces the ability of Nanog to bind to enhancer elements.

We therefore examined the Nanog binding profile at the regulatory regions of several key pluripotency genes using ChIP-seq. There was a clear increase in Nanog binding at several loci following depletion of Oct4 (Figure 4C). Interestingly, expression of these genes is decreased following induced Oct4 degradation despite the increased Nanog binding (Figure 2B and Figure S2B-C). Of particular note, the Klf4 distal enhancer element showed greater Nanog signal following loss of Oct4, yet reduced mRNA and enhancer RNA expression, and H3K27 acetylation (Figure 2B, Figure S2B-C, Figure 3A-B and Figure S3B-C). This further highlights that the presence of Oct4 appears to be absolutely required for the expression of certain pluripotency-associated genes.

### Nanog binding is increased globally, independent of Oct4 co-binding

We further analysed our Nanog ChIP-seq data to examine global changes in Nanog binding immediately following Oct4 ablation. Looking at all Nanog binding sites revealed a global shift towards greater levels of Nanog enrichment (Figure 5A and B). Examining the change in Nanog signal over all peaks, there is a significant overall increase following loss of Oct4 (16608 Nanog peaks; paired t-test of Nanog signal before and after treatment, p < 10^−10^; t-test of log fold change of thresholded peaks vs 4458 sub-threshold (background) peaks, p < 10^−10^). To investigate whether the effect on Nanog binding was specific to sites where Oct4 co-binds, we looked at the complementary subsets of Nanog peaks less than 1kb from an Oct4 peak (3930 sites) and those that do not co-bind with Oct4 (12678 sites), again using peaks called from Oct4 ChIP-seq data from (Marson et al., 2008) (Figure 5B). At Oct4-Nanog co-binding sites there was a detectable increase in Nanog signal; globally this was statistically significant (paired t-test of Nanog signal before and after treatment, p < 10^−10^; t-test of log fold change of thresholded peaks vs 382 sub-threshold (background) peaks, p < 10^−10^), though surprisingly the magnitude of the increase was reduced compared to all Nanog peaks. However, Nanog peaks that do not overlap with Oct4 displayed a more dramatic increase in Nanog binding (12678 peaks; paired t-test of Nanog signal before and after treatment, p < 10^−10^; t-test of log fold change of thresholded peaks vs 4076 sub-threshold (background) peaks, p < 10^−10^), suggesting that the reduced Nanog binding in the presence of Oct4 is not directly due to physical occlusion of Nanog binding sites by Oct4 or due to changes in local chromatin structure in the presence/absence of Oct4. One possible explanation for this global effect is an increase in the stability of Nanog protein following loss of Oct4; however, it has previously shown that Nanog displays increased stability in the presence of Oct4 (Muñoz Descalzo et al., 2013). In keeping with this, we determined the half-life of Nanog with or without addition of IAA which revealed a decrease from ∼ 2.3 hours in the presence of Oct4 to ∼ 1.4 hours in the absence of Oct4 (Figure S5A). This suggests that stability is not the cause of the increased genomic occupancy of Nanog that we observed.

**Figure 5.**
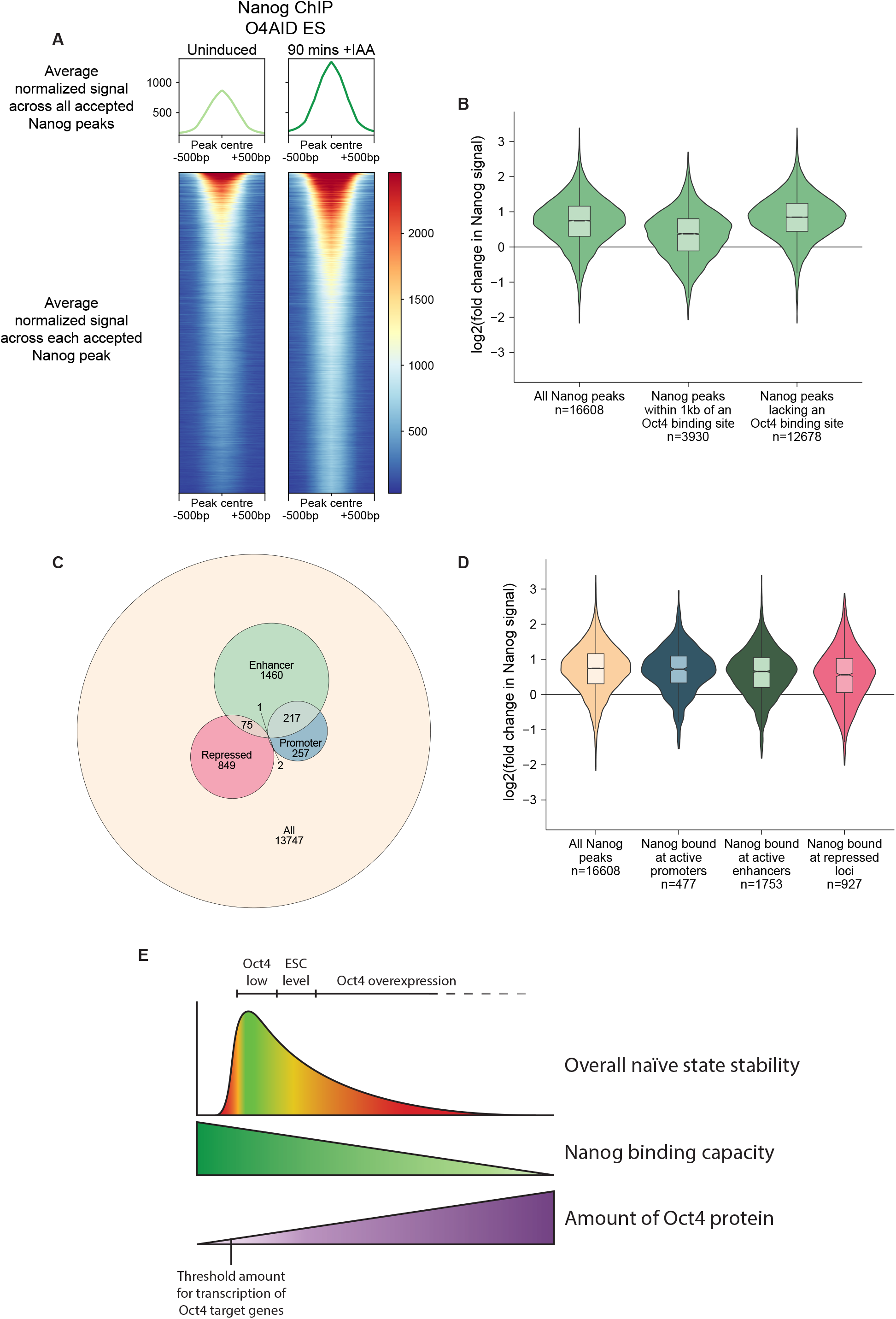
Nanog binding is increased globally, independent of Oct4 co-binding. A) Summary distribution (top) and heatmap (bottom) of Nanog signal (sum of two IPs, normalized to library size) at centred at the summit of all Nanog binding sites across the genome, before and 1.5 hours after addition of IAA. B) Violin and box plot of log2 of the fold change in the average normalized Nanog signal at each Nanog peak in the genome (n=16608) before and after addition of IAA, further broken down into loci within (n=3930) or further than (n=12678) 1kb of Oct4 binding sites generated from ChIP-seq data from Marson et al. 2008. Boxes show the median value and extend to the 25^th^ and 75^th^ quartiles, and whiskers extend to 1.5 times the interquartile range. C) Euler plot showing the number of Nanog peaks assigned to various chromatin environments, and the overlap between assignations. Numbers indicate the number of peaks uniquely in that section of the diagram. D) Violin and box plot of log2 of the fold change in the average normalized Nanog signal at each Nanog peak in the genome before and 1.5 hours after addition of IAA, further broken down into non-exclusive chromatin environments, as indicated in C. Boxes show the median value and extend to the 25^th^ and 75^th^ quartiles, and whiskers extend to 1.5 times the interquartile range. E) Model indicating the proposed relationship between the quantity of Oct4 protein and the capacity for Nanog to bind to the genome and consequently the ability of cells to maintain a naïve identity.

We additionally examined the effect of chromatin context on the change in Nanog binding following loss of Oct4. We used publicly available ChIP-seq data to classify 2861 Nanog peaks as enhancer-promoter- and/or repressive mark associated. As expected, the majority of these Nanog peaks were associated with enhancers (Figure 5C). In all cases, there was a significant increase in Nanog signal (Figure 5D; paired t-test, p < 10^−10^) and the mean increase in signal was comparable, suggesting that the chromatin context has little impact on the ability of Oct4 to restrict Nanog binding under normal conditions.

## Discussion

It has been shown that Oct4 is dispensable for initial generation of the blastocyst structure, but required for segregation of the inner cell mass into naïve epiblast and primitive endoderm (Le Bin et al., 2014). Oct4 is also essential for maintaining naïve pluripotency in vitro (Niwa et al., 2000). Despite this, the manner in which Oct4 is required for these processes is unclear; no overexpression or knockout mutants that rescue the Oct4 knockout phenotype have been identified (Hall et al., 2009; Matoba et al., 2006; Niwa et al., 2000). In order to infer how Oct4 mediates self-renewal, we wanted to examine initial transcriptional changes in response to Oct4 depletion. It is notable that similar experiments have been performed before (Hall et al., 2009; Matoba et al., 2006) and have failed to yield an essential role for Oct4. However, it is known that cells exhibiting a low level of Oct4 can sustain expression of pluripotency markers under mild differentiation conditions (Karwacki-Neisius et al., 2013; Radzisheuskaya et al., 2013), and we wondered if transcription of such factors was being artificially maintained by the gradually reducing level of Oct4 protein in conventional depletion systems. The half-life of Oct4 is sometimes reported to be very long (Lin et al., 2012; Wei et al., 2007); it took a full day for Oct4 to be fully removed from Oct4^F/-^ cells used in this work (Figure 1A-B), and published work shows more than 10 hours for complete Oct4 depletion using a TetOFF system (Hall et al., 2009; Niwa et al., 2000). This is a significant amount of time, especially since transcriptional changes are already observed within this window. In order to avoid this confounding factor, we established new cell lines utilising rapid depletion of Oct4 at the protein level (Figure 1C-H). It has previously been reported that the auxin-inducible degron can reduce the half-life of tagged proteins to ∼20 minutes in mammalian systems (Holland et al., 2012; Nishimura et al., 2009), and indeed we found that tagged Oct4 protein was greatly reduced within half an hour fully depleted within 1.5-2 hours (Figure 1E&H). After ensuring that both the AID domain and Oct4 were unaltered in their function in this fusion protein (Figure 1D&G and Figure S1B-C), we proceeded to re-examine the immediate effects of loss of Oct4. Since depletion of Oct4 was so rapid, we were able to examine the effect of loss of Oct4 prior to changes in the protein level of other pluripotency transcription factors. Crucially, the high temporal resolution afforded by such a rapid depletion system reveals two phases of transcriptional change following loss of Oct, unlike following genetic ablation (Figure 2). First, RNA levels of all pluripotency factors examined were rapidly decreased following degradation of Oct4 protein. Notably, all the factors tested display Oct4 binding at their enhancer or promoter elements in published ChIP-seq data. This suggests that the presence of Oct4 at these elements may be essential for the expression of a broad range of pluripotency associated genes, explaining why no single factor can rescue the Oct4 knockout phenotype. Only subsequently did we observe upregulation of trophectoderm-associated markers, suggesting that this may be a secondary effect. Therefore, repression of an extraembryonic transcriptional network does not seem to be the primary role of Oct4 in mouse ESCs. In keeping with a primary role in permitting active transcription, we found a global reduction in the level of H3K27ac, associated with active enhancers (Figure 3 and Figure S3). The fact that this loss of active enhancer marks extends beyond pluripotency-associated loci could imply that Oct4 is critical for maintaining the uniquely permissive chromatin environment found in naïve pluripotent cells, beyond simply acting to drive expression of key pluripotency factors.

ChIP-qPCR and ChIP-seq against Nanog protein yielded an interesting and surprising result; in the absence of Oct4, greater levels of Nanog were found bound to the genome (Figures 4 and 5, and Figures S4 and 5). Remarkably, this extended beyond Oct4 co-bound sites, with an increase in Nanog levels observed at many binding sites lacking Oct4 (Figure 5B). Consequently, it seems unlikely that this effect is due to Oct4 competing with Nanog for binding sites, in keeping with previous reports of simultaneous binding of Oct4 and Nanog to regulatory regions as detected by sequential ChIP (Medeiros et al., 2009). A previous study examining the localisation of Nanog and Sox2 24 hours after transcriptional depletion of Oct4 found that Sox2 and Nanog binding was reduced at many Oct4 co-bound sites following silencing of Oct4 (King and Klose, 2017). However, this carries the caveat that such changes may be due to subsequent transcriptional and epigenetic changes downstream of loss of Oct4 rather than direct effects. In particular, reduced Nanog signal was associated with loss of transcription and chromatin accessibility, suggesting that these enhancer elements were no longer active or accessible, perhaps due to the onset of differentiation, by the time of their analysis. In contrast, by using a targeted protein degradation system we are able to study the immediate changes in protein localisation in our system, limiting secondary effects.

It is currently unclear how the global nuclear structure changes following acute depletion of Oct4. Previous work has suggested that Oct4 may act as a pioneer factor, opening chromatin that would otherwise be highly compacted (King and Klose, 2017; Soufi et al., 2015). As a result, removal of Oct4 might result in the eviction of other transcription factors and repression of gene expression, though the increased binding of Nanog to chromatin suggests this may not be the case (Figures 4 and 5). Additionally, there could be significant changes in the chromatin architecture due to altered topological domains in the absence of Oct4. Decommissioning of super enhancers, many of which are bound by Oct4 in ESCs (Whyte et al., 2013), could lead to a reorganisation of the nuclear structure as these elements interact with multiple promoter regions, even over large distances (Novo et al., 2018).

The ability to probe transcriptional responses immediately following total Oct4 depletion, and the capacity to examine changes in the binding of Nanog to chromatin prior to significant secondary effects has allowed us to go some way to explaining past discrepancies in the perceived role of Oct4 in the pluripotency network. As Hall et al. note, Nanog has been described as an Oct4 target gene, and there is significant biochemical data validating that Oct4 binds to the Nanog enhancer region and is required for its expression (Kuroda et al., 2005; Rodda et al., 2005), yet gradual Oct4 depletion appeared to have a limited, even positive impact on Nanog mRNA (Figure 2A, Figure S2) (Hall et al., 2009). We propose that the reason for this is that while Oct4 may be essential for Nanog expression, above a minimal level Oct4 negatively regulates the binding of other transcriptional activators. Thus, as Oct4 levels drop and cells pass through an Oct4-low state, binding of Oct4 to the Nanog enhancer is reduced and other transcription factors can be recruited, buffering Nanog expression levels. Once Oct4 is entirely depleted Nanog expression falls as the locus is no longer able to be transcribed.

In agreement with existing literature, it appears that the half-life of Nanog is increased in the presence of Oct4 (Figure S4A) (Muñoz Descalzo et al., 2013). As a result, increased Nanog stability is not likely to be responsible for the increased genomic occupancy we observed. However, it would be interesting to see what other properties of Nanog protein are altered in the absence of Oct4, particularly whether changes in the Nanog protein interactome or DNA binding affinity could explain the increase in chromatin binding. Differences in post-translational modifications could affect the ability of Nanog to interact with DNA and proteins in this way.

Combining the transcriptional responses to loss of Oct4 with the observed global changes in Nanog ChIP signal have led us to a unifying model for the range of phenotypes associated with different levels of Oct4 protein in ESCs (Figure 5E). In the absence of Oct4 key pluripotency factors cannot be expressed and cells cannot maintain a naïve identity. With low levels of Oct4, a threshold is reached such that these factors are able to be expressed, and efficient binding by Nanog results in robust transcription of the whole naïve network to such an extent that differentiation is compromised. At higher Oct4 levels, such as seen in wild-type ESCs and embryos, Oct4 globally supresses the binding of Nanog resulting in a weaker transcriptional network. This allows cells to undergo differentiation in response to signalling cues. On overexpression of Oct4, it is expected that the binding of Nanog to chromatin will be further reduced, destabilizing the naïve network to the point that it cannot be readily maintained.

In summary, utilizing a rapid protein-level depletion strategy, we identified that the primary transcriptional response to loss of Oct4 is a decrease in the expression of pluripotency-associated genes, and upregulation of trophectoderm factors is a subsequent event. Additionally, we found a global increase in the amount of Nanog associated with the genome in the absence of Oct4, suggesting a mechanism by which wild-type levels of Oct4 ensure that naïve cells retain the capacity to initiate differentiation in response to appropriate signals. Together, these reveal a model that ties together the range of phenotypes associated with differing levels of Oct4, concisely explaining how different levels of this factor result in seemingly contradictory cell behaviours.

## Experimental procedures

### Cell culture

Mouse ESCs were cultured in 2iL conditions as previously described. Briefly, cells were maintained in N2B27 (1:1 DMEM/F-12 and Neurobasal, 2mM L-glutamine, 1x penicillin-streptomycin, 0.1mM 2-mercaptoethanol, 1% B27, 0.5% N2) supplemented with 3μM CHIR99021, 1μM PD0325901 and 20ng/ml mLIF. Where stated cells were transitioned into SL conditions, consisting of GMEM without L-glutamine, 10% fetal bovine serum, 1x non-essential amino acids, 1mM sodium pyruvate, 2mM L-glutamine, 1x penicillin-streptomycin, 0.1mM 2-mercaptoethanol, and 20ng/ml mLIF. ES and iPS cells were maintained on gelatin-coated tissue culture plastic, NSCs were maintained on laminin-coated tissue culture plastic. Cells were passaged every 2-4 days using accutase as required. Where described, media was supplemented with 500nM 4-OHT, 1μg/ml doxycycline, and/or 500μM indole-3-acetic acid.

### Cell Lines

Oct4^F/-^ ESCs (Oct4^F/β-geo^) were previously derived from a cross between Oct4^+/β-geo^ and Oct4^F/F^ mice. O4AID ESCs were generated by transfecting ZHBTc4 ESCs (Niwa et al., 2000) with pPB-CAG-Oct4AID-PGK-Hph and pPB-CAG-Tir1-IRES-Bsd with pPBase to achieve efficient integration, and maintained in the presence of Doxycycline to silence the TetOFF-Oct4 transgene. O4AID iPSCs were generated from Oct4-/- NSCs (Radzisheuskaya et al., 2013) nucleofected with pPB-CAG-Oct4AID-PGK-Hph and pPBase to achieve efficient integration and reprogrammed; equivalent wild-type Oct4 control cells were generated by nucleofection with pPB-CAG-Oct4-PGK-Hph and pPBase. The generated O4AID iPSCs were transfected with pCAG-Tir1-IRES-bsd and clonal lines were assessed for functional transgene integration.

### Reprogramming

PLAT-E cells were transfected with pMXs-Oct4, pMXs-Klf4 or pMXs-cMyc using FuGENE 6 reagent to produce retroviral particles. Medium was changed the following day and after 48 hours supernatant containing retroviral particles was collected. Media were filtered and mixed together in equal ratio, then polybrene was added to a final concentration of 4μg/ml. The polybrene/virus mixture was applied to NSCs. 24 hours later, NSCs were nucleofected with a 1:5 ration of pPBase and either pPB-CAG-Oct4AID-PGK-Hph or pPB-CAG-Oct4-PGK-Hph using Amaxa Nucleofection Technology and plated in NSC medium for 2 days then switched to SL medium. Medium was then switched to KSR-2iL (GMEM without L-glutamine, 10% KOSR, 1% fetal bovine serum, 1x non-essential amino acids, 1mM sodium pyruvate, 2mM L-glutamine, 1x penicillin-streptomycin, 0.1mM 2-mercaptoethanol, and 20ng/ml mLIF). Selection was applied for expression from the endogenous Oct4 locus on the ninth day in KSR-2iL. Once colonies had expanded they were passaged into 2iL conditions.

### qRT-PCR and RNA-seq

Total RNA was isolated from cultured cells using the RNeasy Mini Kit (Qiagen) according to the manufacturer’s instructions including on-column DNaseI digest. For RT-qPCR, RNA was reverse-transcribed using SuperScript III First-Strand Synthesis SuperMix for RT-qPCR and reactions were performed on a StepOnePlus Real time PCR system with recommending settings using TaqMan Fast Universal PCR Mix or Fast SYBR Green Master Mix.

For high throughput RNA sequencing, RNA integrity was assessed on a Qubit Fluorometer (ThermoFisher Scientific) and Agilent Bioanalyzer Nano Chips (Agilent Technologies). Depletion of ribosomal RNA was performed on 2-5μg of total RNA using the Ribo-Zero rRNA Removal Kit (Illumina) and libraries were produced from 10-100ng of ribosomal-depleted RNA using NextFlex Rapid Directional RNA-seq Kit (Bio Scientific) with 12 cycles of PCR amplification. Libraries were pooled in equimolar quantities and sequenced on the HiSeq4000 platform (Illumina) at CRUK. Library preparation was performed by the W-MRC CSCI genomics facility. Reads were aligned to mouse genome reference GRCm38/mm10 with TopHat2 (Kim et al., 2013) v2.1.0 (https://ccb.jhu.edu/software/tophat) using default parameters for paired end reads. Gene-wise counts were generated using featureCounts (Liao et al., 2014) based on annotation from Ensembl GRCm38.86 release. Transcript counts were TMM normalized and differentially expressed genes were called with q>0.9 using NOISeq-sim from NOISeq (Tarazona et al., 2011, 2015) version 2.30.0. For all NOISeq analyses, the following parameters were used: pnr=0.2, nss=5, v=0.02; seed ‘321’ was used for each analysis.

### Western Blot

Cells were lysed in RIPA buffer (Sigma) containing Complete-ULTRA protease-inhibitor and PhoStop phosphatase-inhibitor cocktails (Roche), and sonicated with Bioruptor200 (Diagenode) at high frequency, alternating 30 s on/off for 3 min. SDS-PAGE electrophoresis was performed using Bolt 10% Bis-Tris Plus gels (ThermoFisher) in a Novex MiniCell (ThermoFisher). Protein transfer was performed using the semi-dry iBlot2 system (ThermoFisher) and iBlot Transfer Stacks (ThermoFisher). Primary antibodies were as follows: Oct4 (Monoclonal rabbit anti-Oct4, Cell Signalling Technology, #83932), Nanog (Monoclonal rat anti-Nanog, eBioscience, #14-5761-80), Sox2 (Monoclonal rat anti-Sox2, eBioscience, #14-9811-80), alpha-tubulin (Monoclonal mouse anti-alpha-Tubulin, Abcam, #ab7291). Detection was achieved using HRP-linked secondary antibodies diluted 1:10000 against the appropriate species (GE Healthcare) and ECL Plus Western Blotting Detection System (GE Healthcare).

### Protein half-life analysis

Cells were treated with cycloheximide with or without addition of IAA, and harvested at hourly intervals. Western blot analysis of Nanog protein levels were performed, and a relative standard curve was included using serial dilutions of uninduced sample. Western blots were quantitated using Image Studio Lite Quantification Software.

### ChIP-qPCR and ChIP-seq

ChIP was performed as previously described. Briefly, 10×10^6^ cells per IP for transcription factors or 5×10^6^ cells per IP for histone modifications were fixed for 10 min in 1% formaldehyde. Fixation was halted by addition of an excess of glycine. Cells were washed with ice-cold PBS. Nuclei were isolated: Cells were incubation with Lysis Buffer 1 (50 mM HEPES pH 7.5, 140 mM NaCl,1 mM EDTA, 10% Glycerol, 0.5% NP40, 0.25% Tx100) at 4 °C for 10 mins, pelleted, then incubated with Lysis Buffer 2 (10 mM Tris pH 8.0, 200 mM NaCl, 1 mM EDTA, 0.5 mM EGTA) at 4 °C for 10 mins. Nuclei were pelleted then resuspended in Shearing Buffer (50 mM Tris pH 8.0, 1% SDS, 10 mM EDTA) and sonicated to obtain an average DNA fragment size of 500 base pairs. Lysates were diluted 1:10 in dilution buffer (50 mM Tris–HCl at pH 8.0, 167 mM NaCl, 1.1% Triton X-100 and 0.11% sodium deoxycholate) and pre-cleared for 2 h at 4 °C with Dynabeads Protein G magnetic beads (Invitrogen) that were pre-incubated with isotypic IgG antibody. Supernatent was collected, a 10% input sample was taken for relative quantitation, and the chromatin was the incubated overnight at 4 °C with 2 μg of antibody or 2 μg of isotypic IgG control. Lysates were then incubated for 1 h at 4 °C with BSA-blocked Dynabeads Protein G magnetic beads, and the beads were washed twice in wash buffer 1 (50 mM Tris pH 8.0, 0.1% SDS, 0.1% Sodium deoxycholate, 1% Tx100, 150 mM NaCl, 1 mM EDTA, 0.5 mM EGTA), once in Wash Buffer 2 (50 mM Tris pH 8.0, 0.1% SDS, 0.1% sodium deoxycholate, 1% Tx100, 500 mM NaCl, 1 mM EDTA, 0.5 mM EGTA), once in Wash Buffer 3 (50 mM Tris pH 8.0, 250 mM LiCl, 0.5% Sodium deoxycholate, 0.5% NP40, 1 mM EDTA, 0.5 mM EGTA) and twice in wash buffer 4 (50 mM Tris pH 8.0, 10 mM EDTA and 5 mM EGTA). Chromatin was eluted twice for a total of 30 min at 65 °C in elution buffer (1% SDS and 0.1 M NaHCO_3_). NaCl was added to samples and inputs to a final concentration of 200 mM, before incubating overnight at 65 °C to reverse the crosslinking and DNA was purified using the QIAquick PCR Purification kit (Qiagen).

Chromatin was analysed by qPCR using a StepOnePlus Real time PCR system with recommending settings and Fast SYBR Green Master Mix. Additionally, next generation sequencing libraries were prepared using the ThruPLEX DNA-Seq Kit. Libraries were pooled in equimolar quantities and sequenced on the HiSeq4000 platform (Illumina) at CRUK. Library preparation was performed by the W-MRC CSCI genomics facility. Reads were aligned to mouse genome reference GRCm38/mm10 using Bowtie 2 (Langmead and Salzberg, 2012) version 2.3.4.3 with default parameters and deduplicated using SAMtools rmdup. Normalized enrichment tracks were generated with the MACS2 (Liu, 2014) callpeak function using the –SPMR flag. H3K27ac peaks were called using MACS2 callpeak function on pooled data from uninduced samples. For analysis of signal intensity, replicates were pooled and RPKM normalized using DiffBind (Stark and Brown, 2017). Peaks with RPKM < 0.5 in both uninduced and induced samples were considered to be background and excluded from further analysis. Subsets of peaks located within 1kb of Oct4 peaks were determined by analysis of published ChIP-seq data; reads were deduplicated and aligned as above, and peaks were called using MACS2 callpeak function with whole cell extract used as control samples. Nanog peaks were called using MACS2 callpeak function on pooled ChIP-seq data from both uninduced and IAA treated samples. For analysis of Nanog signal at binding sites, replicates of treated and untreated samples were pooled and RPKM normalized using deepTools2 (Ramírez et al., 2016) bamCoverage and blacklisted regions from the ENCODE project consortium were excluded, then a heatmap and summary plot were generated using deepTools2 computeMatrix and plotHeatmap tools. For pairwise analysis of Nanog binding at peaks called from pooled data from both uninduced and IAA treated samples, centred on the peak summit and extending a total of 350bp, replicates were pooled and RPKM normalized using DiffBind. Peaks with an RPKM < 0.9 in both uninduced and induced samples were considered to be background and excluded from further analysis. Subsets of peaks corresponding to active promoters, active enhancers and repressed regions were assigned using published ChIP-seq peaksets from ENSEMBL, using the following criteria: active promoters, within 1kb of PolII and either overlapping H3K4me1 or H3K4me3; active enhancer, within 1kb of P300 and overlapping H3K27ac; repressed region, overlapping either H3K9me3 or H3K27me3.

### Public datasets used

| Dataset | Source | Analysis |
| --- | --- | --- |
| Bruce4 ES EP300 ChIP-seq peakset | Ensembl (release 100) | SWEmbl |
| Bruce4 ES PolII ChIP-seq peakset | Ensembl (release 100) | SWEmbl |
| Bruce4 ES H3K27ac ChIP-seq peakset | Ensembl (release 100) | SWEmbl |
| Bruce4 ES H3K27me3 ChIP-seq peakset | Ensembl (release 100) | CCAT |
| Bruce4 ES H3K4me1 ChIP-seq peakset | Ensembl (release 100) | CCAT |
| Bruce4 ES H3K4me3 ChIP-seq peakset | Ensembl (release 100) | SWEmbl |
| Bruce4 ES H3K9me3 ChIP-seq peakset | Ensembl (release 100) | CCAT |
| V6.5 ES Oct4 ChIP-seq data | Marson et al. 2008, GEO accession GSM307137 | See above |
| V6.5 ES whole cell extract data | Marson et al. 2008, GEO accessions GSM307154 and GSM307155 | See above |

### Antibodies

| Target | Source | Catalogue number |
| --- | --- | --- |
| Polyclonal goat anti-Oct4 | Santa Cruz Biotechnology | sc-8628 |
| Monoclonal mouse anti-alpha-Tubulin | Abcam | ab7291 |
| Polyclonal rabbit anti-H3K27ac | Abcam | ab4729 |
| Monoclonal rat anti-Nanog | eBioscience | 14-5761-80 |
| Monoclonal rat anti-Sox2 | eBioscience | 14-9811-80 |

### qPCR primers and probesets

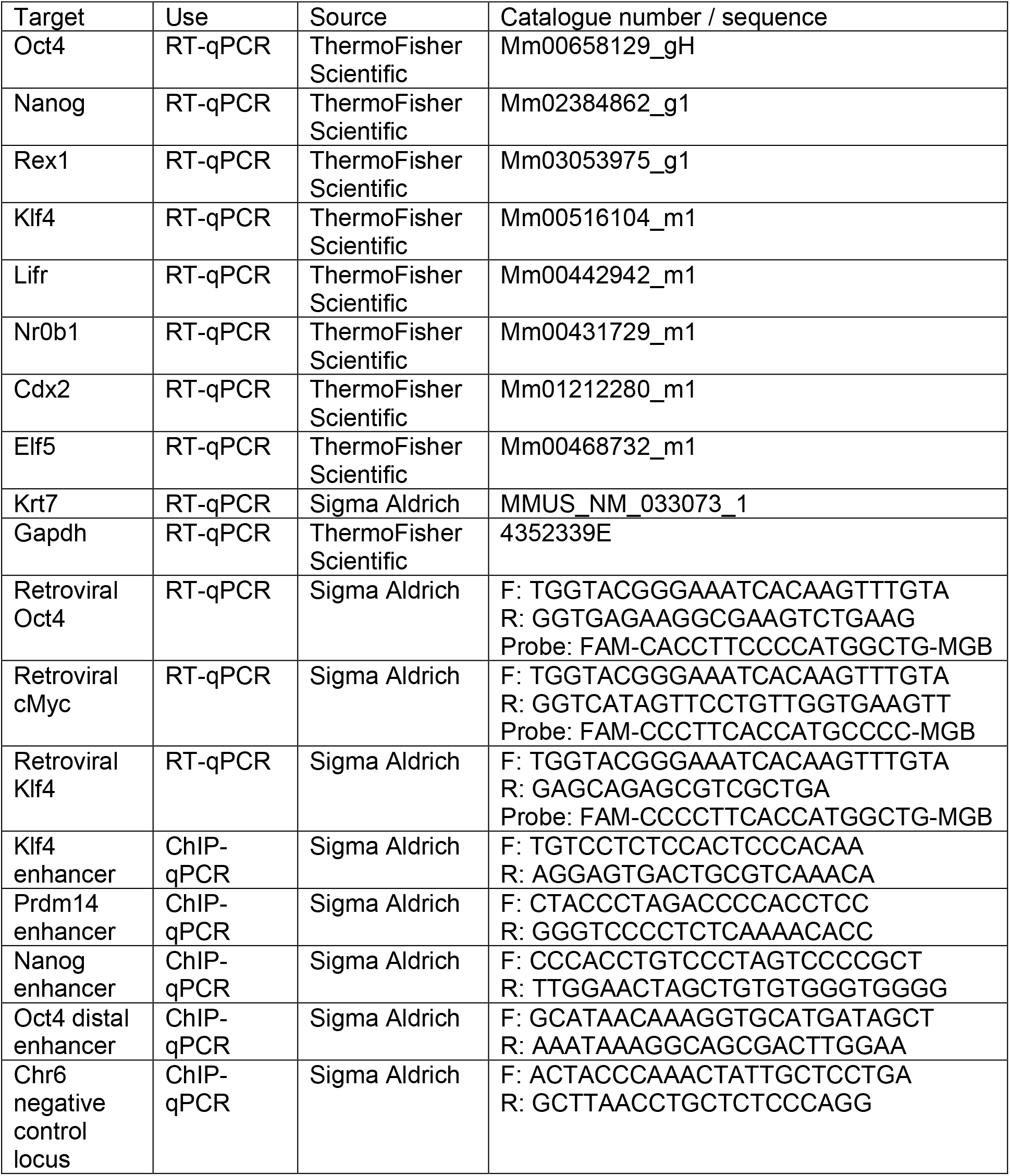

## Acknowledgments

We thank S. Lees, M. Paramor, M.A. Barber, and P. Humphreys for specialist technical support. We are grateful to A. Radzisheuskaya for technical assistance and Y. Costa and I. Stockwell for critical reading of the manuscript. This study was supported by a Wellcome Trust Fellowship (WT101861) to J.C.R.S and by BBSRC and MRC research grants, BB/R018588/1 and MR/R017735/1 respectively. The authors gratefully acknowledge core support from the Wellcome-MRC Cambridge Stem Cell Institute.

## Author Contributions

L.E.B. and J.C.R.S. conceived the study. L.E.B. designed and performed experiments and bioinformatics analysis, analysed the data, supervised the study and wrote the manuscript. M.R.P.A. performed experiments. J.C.R.S. supervised the study and wrote and approved the manuscript.

