## Supplemental Figures for "Auxin-degron system identifies immediate mechanisms of Oct4"

Figure S1. Relating to Figure 1.

A) Cells before and after conventional tamoxifen-induced CreER driven genetic ablation of Oct4. Uninduced cells grow as compact, round, domed colonies. 4 days after application of tamoxifen, cells have attained an enlarged, flattened morphology. Scalebar – 100µm. B) $Oct4^{-/-}$ Tet-OFF Oct4 ESCs (left) and O4AID ESCs share similar morphology, with mostly compact, round, domed colonies. Scalebar – 50µm. C) Oct4-/- NSCs (left) were reprogrammed in the presence of either wild-type Oct4 or Oct4-AID fusion protein. In both cases, similar round, domed colonies were formed (right). Scalebar – 100µm.

**Figure S1**

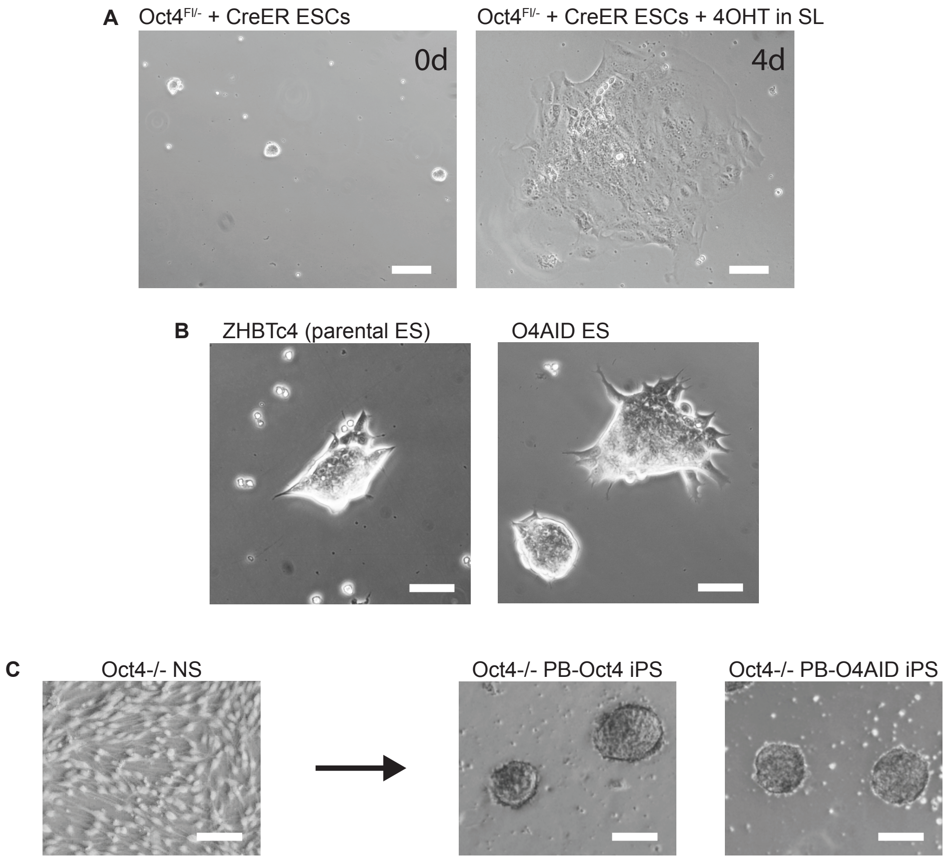
A
Oct4Fl/- + CreER ESCs
Oct4Fl/- + CreER ESCs + 4OHT in SL
0d
4d
B
ZHBTc4 (parental ES)
O4AID ES
C
Oct4-/- NS
Oct4-/- PB-Oct4 iPS
Oct4-/- PB-O4AID iPS

Figure S2. Relating to Figure 2.

RT-qPCR analysis of pluripotency (top) and trophectoderm (bottom) associated genes in Oct4F/- CreER ESCs (A), O4AID ESCs (B), and O4AID iPSCs (C), in a timecourse following induced depletion of Oct4 with 4OHT or IAA. RT-qPCR data represent the mean ± SD of three technical replicates.

#### Figure S2

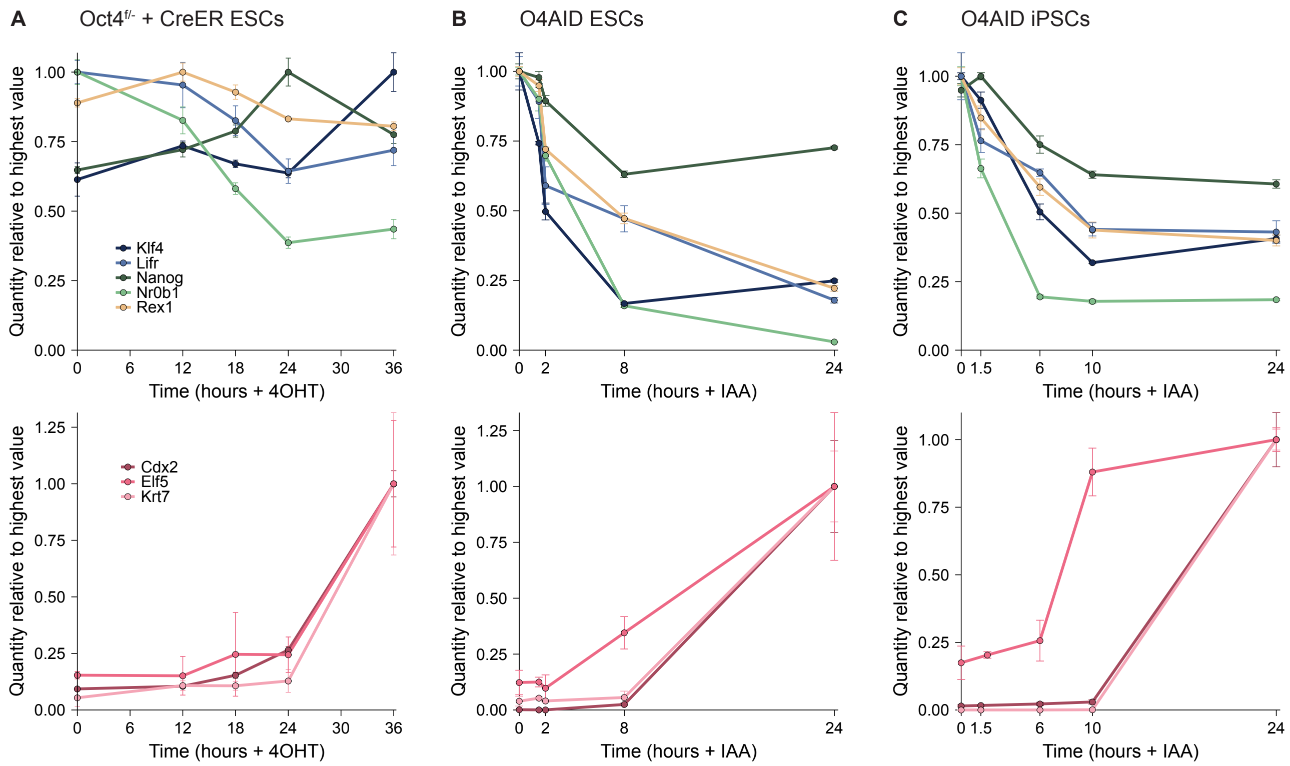

A
Oct4f/- + CreER ESCs
B
O4AID ESCs
C
O4AID iPSCs
Quantity relative to highest value
Klf4
Lifr
Nanog
Nr0b1
Rex1
Cdx2
Elf5
Krt7
Time (hours + 4OHT)
Time (hours + IAA)

Figure S3. Relating to Figure 3.

A) ChIP-seq analysis of H3K27ac at the Tet1 enhancer (top) and the Tet2 enhancer (bottom) in O4AID ESCs before and 1.5 hours after addition of IAA. Genomic coordinates refer to the GRCm38/mm10 genome assembly, and gene intron/exon annotations are taken from Ensembl. Oct4 binding sites generated from ChIP-seq data from Marson et al. 2008 indicated in purple. B&C) ChIP-qPCR analysis of H3K27ac at B) the Klf4 distal enhancer and C) the Prdm14 distal enhancer in O4AID ESCs (left) and O4AID iPSCs (right) before and 1.5 hours after addition of IAA. RT-qPCR data represent the mean ± SD of three technical replicates. ChIP-qPCR data represent the mean ± SD of three IPs for H3K27ac, and the mean of three technical replicates of a single IP for IgG. ND = not detected. * $P<0.05$, *** $P<0.001$ (Student's t test, two-tailed, unpaired, assuming equal variance). D) Violin and box plot showing H3K27ac signal before and 1.5 hours after addition of IAA at all detected H3K27ac peaks. Peaks with signal below the indicated threshold both before and after induction were consider to be background and were excluded from further analysis. E) Violin plot showing log2-fold change in H3K27ac signal between uninduced and 1.5 hour IAA treated O4AID ESCs at the peaks below the threshold indicated in D).

#### Figure S3

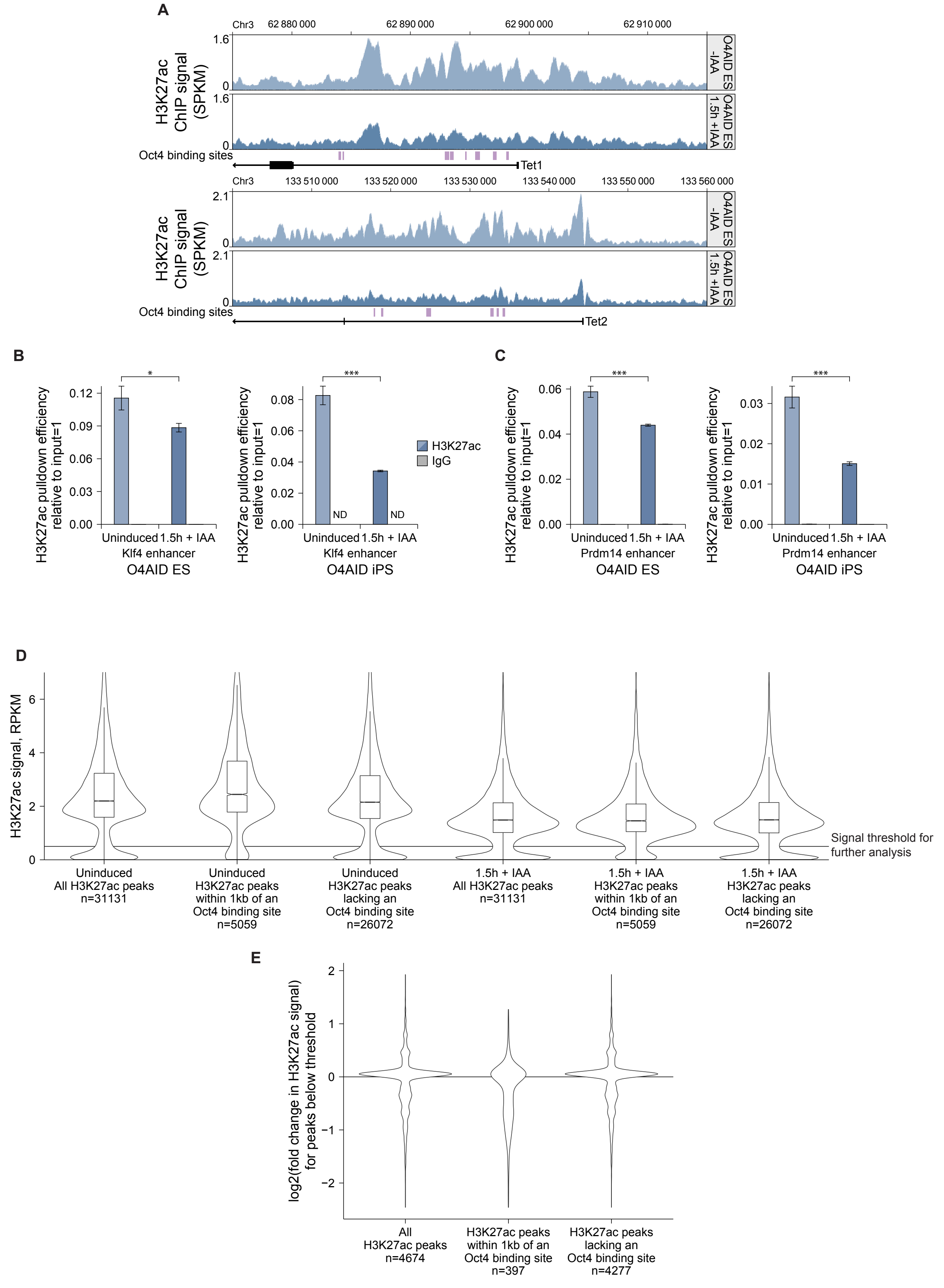

A
Chr3
62 880 000
62 890 000
62 900 000
62 910 000
H3K27ac ChIP signal (SPKM)
O4AID ES -IAA
O4AID ES 1.5h +IAA
Oct4 binding sites
Tet1
133 510 000
133 520 000
133 530 000
133 540 000
133 550 000
133 560 000
Tet2
B
H3K27ac pulldown efficiency relative to input=1
Uninduced
1.5h + IAA
Klf4 enhancer
O4AID ES
O4AID iPS
ND
H3K27ac
IgG
C
Prdm14 enhancer
D
H3K27ac signal, RPKM
Signal threshold for further analysis
Uninduced All H3K27ac peaks n=31131
Uninduced H3K27ac peaks within 1kb of an Oct4 binding site n=5059
Uninduced H3K27ac peaks lacking an Oct4 binding site n=26072
1.5h + IAA All H3K27ac peaks n=31131
1.5h + IAA H3K27ac peaks within 1kb of an Oct4 binding site n=5059
1.5h + IAA H3K27ac peaks lacking an Oct4 binding site n=26072
E
log2(fold change in H3K27ac signal) for peaks below threshold
All H3K27ac peaks n=4674
H3K27ac peaks within 1kb of an Oct4 binding site n=397
H3K27ac peaks lacking an Oct4 binding site n=4277

Figure S4. Relating to Figure 4.

Analysis of Nanog binding to pluripotent regulatory sequences in O4AID iPSCs. A) Protein level of Oct4 and Nanog (Western blot) in O4AID iPSCs before and 2 hours after addition of IAA with α-tubulin as a loading control. B) ChIP qPCR following pulldown of Nanog or using normal IgG negative control at Nanog binding sites or a negative control locus in O4AID iPSCs. Nanog pulldown represents mean ± SD of three IPs; IgG pulldown represents the mean of three technical replicates of a single IP.

#### Figure S4

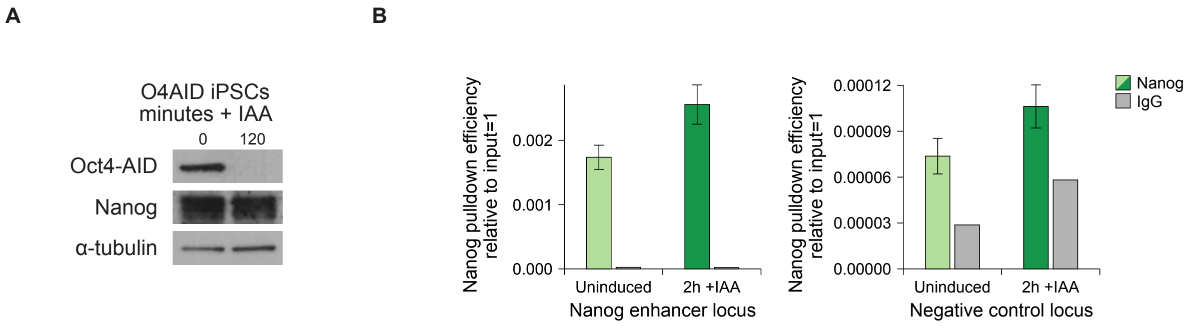
A
B
O4AID iPSCs
minutes + IAA
0
120
Oct4-AID
Nanog
α-tubulin
Nanog pulldown efficiency relative to input=1
0.000
0.001
0.002
Uninduced
2h +IAA
Nanog enhancer locus
0.00000
0.00003
0.00006
0.00009
0.00012
Negative control locus
Nanog
IgG

Figure S5. Relating to Figure 5.

A&B) Analysis of Nanog protein half-life in O4AID ES with and without Oct4 depletion. A) Western blot analysis of Nanog protein level in a timecourse following cycloheximide treatment, with or without addition of IAA. B) Plot of natural log of Nanog protein quantity against time used to calculate the decay constant, $\lambda$, from the equation $\ln(x(t)) = \ln(x_0) + \lambda t$. From this, the protein half-life is calculated as $t_{1/2} = \ln(2)/\lambda$. C) Violin and box plot showing Nanog signal before and 1.5 hours after addition of IAA at all detected Nanog peaks. Peaks with signal below the indicated threshold both before and after induction were consider to be background and were excluded from further analysis. E) Violin plot showing log2-fold change in Nanog signal between uninduced and 1.5 hour IAA treated O4AID ESCs at the peaks below the threshold indicated in C).

### Figure S4

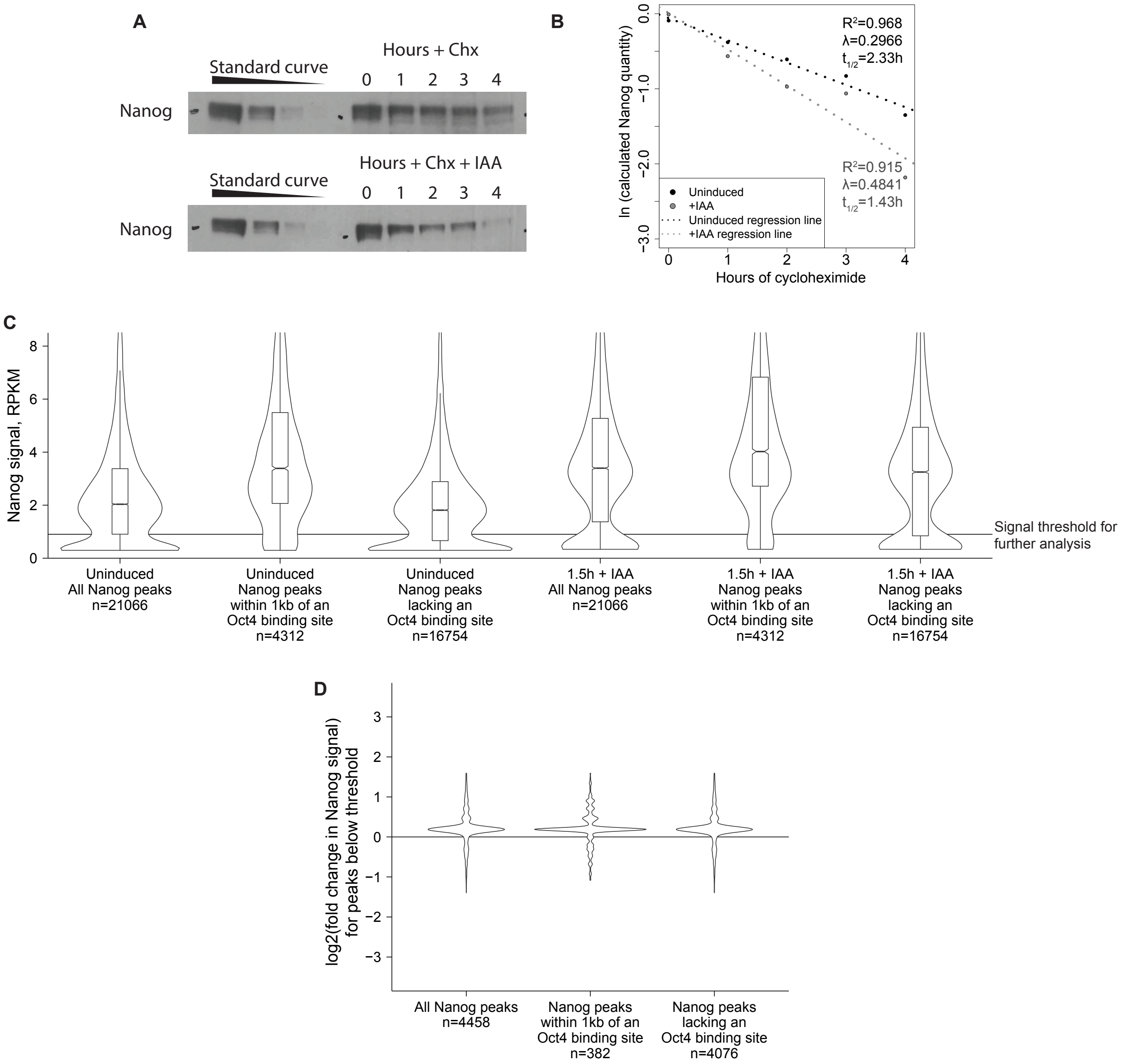

A
Standard curve
Hours + Chx
0 1 2 3 4
Nanog
Standard curve
Hours + Chx + IAA
0 1 2 3 4
Nanog
B
ln (calculated Nanog quantity)
Hours of cycloheximide
$R^2$=0.968
λ=0.2966
$t_{1/2}$=2.33h
$R^2$=0.915
λ=0.4841
$t_{1/2}$=1.43h
Uninduced
+IAA
Uninduced regression line
+IAA regression line
C
Nanog signal, RPKM
Signal threshold for further analysis
Uninduced All Nanog peaks n=21066
Uninduced Nanog peaks within 1kb of an Oct4 binding site n=4312
Uninduced Nanog peaks lacking an Oct4 binding site n=16754
1.5h + IAA All Nanog peaks n=21066
1.5h + IAA Nanog peaks within 1kb of an Oct4 binding site n=4312
1.5h + IAA Nanog peaks lacking an Oct4 binding site n=16754
D
log2(fold change in Nanog signal) for peaks below threshold
All Nanog peaks n=4458
Nanog peaks within 1kb of an Oct4 binding site n=382
Nanog peaks lacking an Oct4 binding site n=4076
